# Extracellular matrix topography drives adrenergic to mesenchymal transition in neuroblastoma

**DOI:** 10.1101/2023.10.10.561780

**Authors:** Antonios Chronopoulos, Chandra Vemula, Ivan Chavez, Rebekah Kennedy, Shahab Asgharzadeh, JinSeok Park

## Abstract

Neuroblastoma (NB), the most common extracranial solid tumor in children, exhibits significant intra-tumoral heterogeneity with two interconvertible identities: adrenergic (ADRN) and mesenchymal (MES). MES cells exhibit phenotypes associated with metastasis and are enriched in relapse NB compared to ADRN. Thus, reprogramming from ADRN to MES may determine inferior NB outcomes, which needs better elucidation. Extracellular matrix (ECM) is an essential tumor microenvironment (TME) component that provides physical support as a scaffold and delivers mechanical cues. We demonstrate that high-risk NB has more topographically aligned ECM fibers than low-risk NB. Using nano-fabricated biomaterials mimicking ECM alignment, we reveal that ECM topography drives ADRN-MES reprogramming by enhancing cell-ECM interactions. This transition involves epigenetic and transcriptional changes, accompanied by enhanced phenotypic features of MES. Also, we uncover that ECM-driven reprogramming relies on the Rho-associated kinase pathway. Overall, ECM-driven ADRN-MES reprogramming provides insight into TME-targeted therapeutic strategies for suppressing MES and improving NB outcomes.

## Main text

Neuroblastoma (NB) is the most common extracranial pediatric solid tumor, responsible for 15% of cancer-related deaths in children. It exhibits remarkable heterogeneity composed of two interconvertible cell identities: adrenergic (ADRN) and mesenchymal (MES) (1–4). Core regulatory circuitries (CRCs), a set of lineage transcription factors (TFs), define the identity by modulating the underlying epigenetic and transcriptional landscape (1, 3, 5). Their distinct phenotypes characterized by their underlying molecular landscape are closely related to tumor progression, treatment resistance, and disease outcome (1, 3, 4). ADRN cells exhibit features similar to normal sympathetic neurons and have a more differentiated, less aggressive behavior. In contrast, the less differentiated MES, reminiscent of mesenchymal and stem-like features, is associated with invasiveness, metastasis, and therapy resistance(6). Recent research has illuminated the reversible nature of ADRN and MES states(4). Lineage TF modulation can lead to switching via remodeling epigenetic and transcriptional landscapes, mimicking the natural interconversion(7). The transition toward MES influences NB progression and treatment responses, leading to inferior NB outcomes. Thus, we need better elucidation of how NB modulates the switch between MES and ADRN.

The impact of the tumor microenvironment (TME) on NB progression has gradually gained attention (8, 9). However, the role of the extracellular matrix, an essential TME component defining the mechanical properties of tumor stroma, has been underappreciated. The ECM provides structural support as a scaffold and serves as a signaling platform that influences cell characteristics via mechano-biological regulation(10). Intriguingly, highrisk NB exhibited enriched reticulin fibers composed of ECM molecules in the TME(11). Also, ECM has been suggested to control molecular features related to NB outcomes(12, 13). Although these findings suggest the profound effects of ECM on NB progression, it remains understudied.

Cancer cells are sensitive to ECM mechanical features, such as its stiffness and topography (14–18). Their mechano-sensing induces cells to establish focal adhesions, protein complexes anchoring cells to the ECM substrate and involved in sensing underlying mechanical ECM features further(19). The focal adhesion mediates cytoskeletal reorganization and triggers signaling cascades that influence chromatin remodeling, transcription factor activity, and, ultimately, gene and protein expression(10, 16, 20, 21). Beyond oncogenic alterations, cell-ECM interactions have been shown to influence every hallmark of cancer(22).

Specifically, ECM topography, characterized by patterns of geometrical features, e.g., ridges and grooves, can influence cellular behavior related to cancer progression. Tumor cells exert forces and proteolytic enzymes to reorganize collagen, a major ECM molecule, at the tumor-stroma interface (23). The remodeled collagen results in a distinct matrix topography displaying fibril alignment. The aligned ECM reciprocally feeds back to cancer cells to facilitate local invasion(23). Furthermore, aligned ECM topography can drive chemoresistance in breast cancer by regulating cell signaling and metabolism(24).

Herein, we investigate the effect of ECM topography and cell-ECM interactions on ADRN-MES transition in NB. We used biomaterial substrates with nanogrooved arrays (termed thereafter NGA) as an *in vitro* model of ECM alignment. The NGA recapitulates the structural, dimensional, and topographical alignment of collagen fibrils observed in high-risk NB tumors, thus maintaining high pathophysiological relevance. Additionally, NGA allows robust experimental compatibility with microscopy applications and downstream biochemical assays(25). We have combined our biomaterial strategy with -omics approaches and functional assays to dissect the mechanobiological-mediated changes in the epigenome, transcriptome, and protein expression, to unveil how the biophysical TME affects NB progression.

Our data show that the ECM topography in the TME induces ADRN-MES reprogramming. Its effects on ADRNMES transition involved mechanobiological modulation of the epigenetic state and transcriptional rewiring, which requires Rho-kinase signaling (ROCK) stimulated by cell-ECM interaction. Functionally, the ADRN-MES phenotypic transition displayed elevated resistance to ALK inhibitors and increased invasiveness. The limited number of available MES models compared to ADRN has hampered research efforts into NB heterogeneity(2). In this regard, our biomaterial strategy provides an efficient way to overcome this limitation. Overall, ECM topography-driven ADRN-MES reprogramming sheds light on the TME regulation of NB progression and offers exciting avenues for therapeutic exploration.

### ECM topography enhances transcriptional ADRN-MES reprogramming

To identify different molecular characteristics between MES and ADRN, we conducted gene set enrichment analysis (GSEA) with the transcriptome of MES versus ADRN cell lines. GSEA results revealed significant enrichment of gene sets related to collagen biosynthesis, remodeling, and cell-ECM interactions for the MES subtype (Fig. 1a, Supplementary Fig 1, and Supplementary Table 1). The result suggests that high-risk NB with a conceivably higher subpopulation of MES cells have TME conditions promoting cell-ECM interaction. Indeed, we found that high-risk and relapsed tumors demonstrated highly enriched ECM compared to the low-risk samples through immunohistochemical (IHC) staining with collagen type III, a representative ECM molecule in NB (Fig 1b). Their stromal ECM fibers displayed a more aligned topography and higher staining intensity compared to the haphazard organization in low-risk NB (Fig. 1c and Supplementary Figure 2).

**Figure 1:**
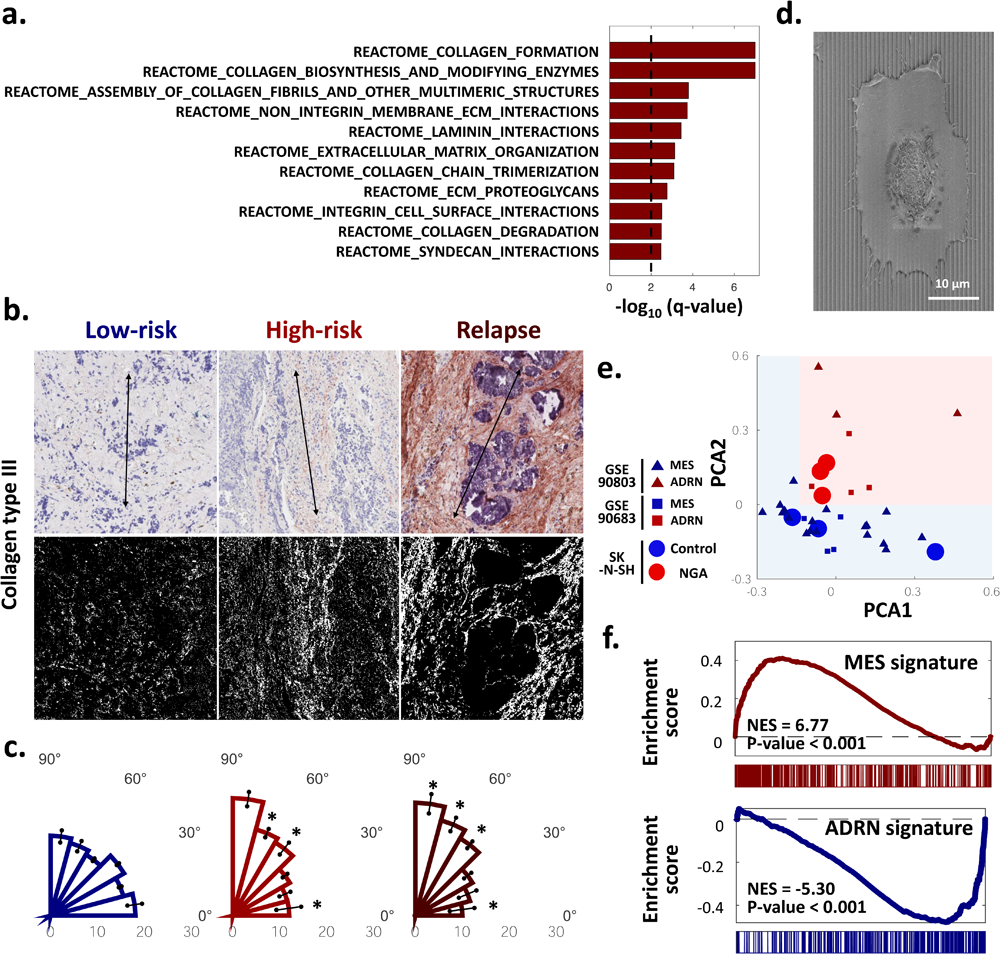
NGA substrates, mimicking aligned ECM topography in high-risk and relapsed NB, induce transcriptional reprogramming from ADRN to MES. **(a).** Gene set enrichment analysis (GSEA) with transcriptome highly expressed in MES compared to ADRN (GSE90683). Reference gene sets are REACTOME. The result suggests that MES exhibits higher enrichment of gene sets related to the synthesis and remodeling of ECM (e.g., collagen and laminin) and cell-ECM interactions. **(b).** Immunohistochemical staining of different-risk NB tissues with collagen type III antibody. **(c).** Quantification of topographical alignment degrees of collagen type III (Col-III). Double-sided arrows in Panel (b) indicate 90°. Statistical difference of the populational ratio of Col-III alignments in the same angle range between low-risk and high-risk/relapse NB. Two-sided t-test. P-value*<0.05, n=3, Error bars are S.E.M **(d).** Scanning electron microscopic image of a cell on NGA. **(e).** Principal component analysis with the transcriptome of MES and ADRN (GSE90803 and GSE90683) and that of SK-N-SH cultured on Col-III pre-coated NGA and control flat substrates. A red-shaded region indicates ADRN-like cells and a blue-shaded region means MES-like cells. **(f).** GSEA with highly expressed genes in SK-N-SH cells cultured on NGA compared to control flat substrate, suggesting their positive enrichment of MES signatures and depletion/negative enrichment of ADRN signatures.

We next ascertained whether their topographical alignment has a causal role in ADRN-MES transition. We used NGA coated with collagen type III to mimic the native ECM topographical alignment in high-risk and relapsed tumors. NGA was fabricated by capillary-force lithography with a biocompatible polymer. Its topographical feature size (arrays of 1μm-size grooves/ridges) approximates the diameter of ECM fibers known to influence cell fate. (Fig. 1d). A collagen type III-coated flat surface of the same polymer without topographical features was used as a control. To evaluate transcriptional reprogramming by ECM topography, we performed RNA-seq with SK-N-SH cells (NB cell line with mixed ADRN/MES subpopulations) plated on NGA and flat substrates. Interestingly, principal component analysis of their transcriptome acquired by RNA-seq revealed clustering of NGA cells in the groups of dominant MES subtypes (Fig. 1e). In contrast, NB cells on control substrate clustered around ADRN cells. Furthermore, GSEA revealed distinctively higher MES signature enrichment in cells plated on NGA (relative to flat control) with concomitant suppression of ADRN signatures (Fig. 1f, Supplementary Table 2). Taken together, these results suggest that the distinctive topography of ECM in high-risk/relapsed NB has a causal role in mechano-biologically rewiring the transcriptional landscape that triggers ADRN-MES transition.

### Cell identity switching towards MES by ECM topography

We evaluated whether the observed transcriptional differences on NGA translate to switching toward MES at the translational level. We performed immunofluorescent (IF) staining against vimentin(3), a cytoskeletal marker indicative of the MES state, and PHOX2B(3), a transcription factor critically involved in the development of sympathetic neurons that has ADRN-specific expression. NB cells (SK-N-SH, CHLA-255, and SK-N-AS) on NGA had higher expression of vimentin and lower levels of PHOX2B relative to flat controls as measured by their respective IF intensity values (Fig. 2a,b and Supplementary Fig. 3a). PRRX1 is a core MES driver that can reprogram the epigenetic and transcriptomic circuitries of ADRN cells towards the MES state(3). We consistently found higher PRRX1 expression along with lower expression level of GATA3, the ADRN marker, on NGA than on control flat surfaces (Fig 2c).

**Figure 2:**
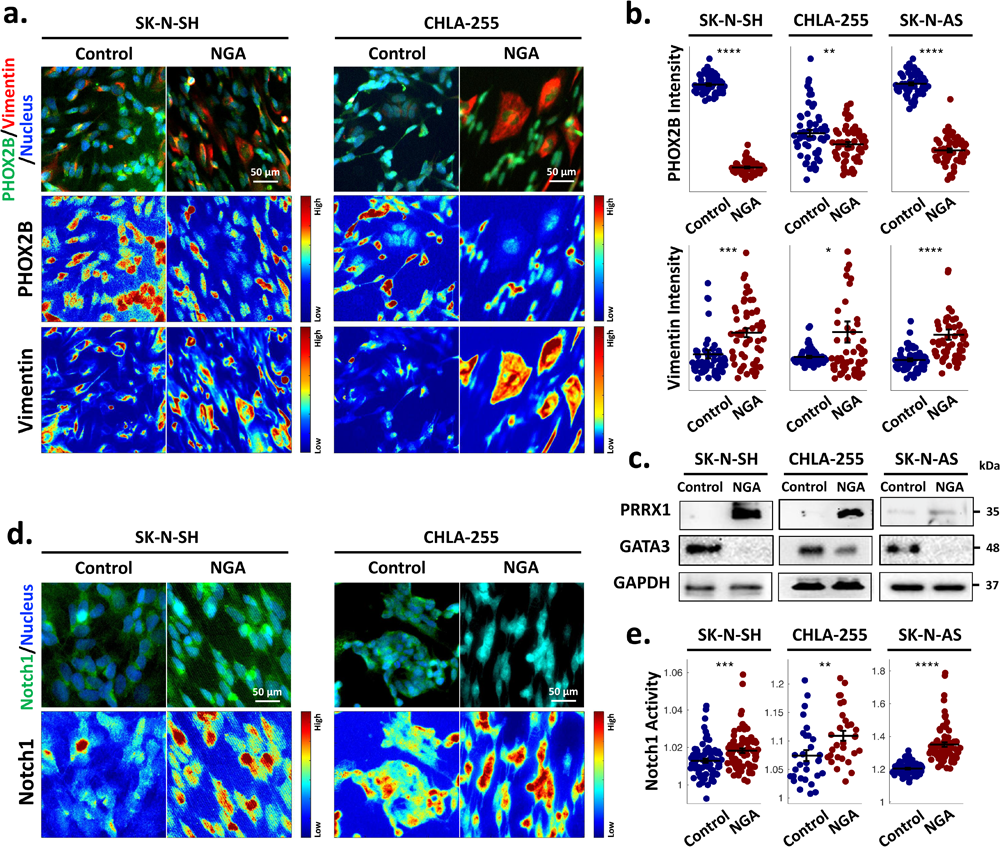
NB cells on NGA exhibit increased expression of MES markers and decreased expression of ADRN markers. **(a).** Immunofluorescent (IF) staining of NB cells on control flat and NGA substrates with antibodies of PHOX2B (ADRN marker) and vimentin (MES marker). Pseudo-color images indicate IF intensity. **(b).** IF intensities of corresponding markers in Panel **A. (c).** Western blotting (WB) of NB cells on flat and NGA substrates with antibodies against PRRX1 (MES marker) and GATA3 (ADRN marker). **(d).** Notch1 IF staining of cells on flat and NGA surfaces. **€.** Notch1 activity of cells on control and NGA substrates, quantified by the ratio of nuclear Notch1 IF intensity to cytoplasmic Notch1 IF intensity. In Panel **B** and **E**, each dot indicates the value of a corresponding cell. Lines indicate mean values of corresponding groups. Error bars are S.E.M. Statistical difference of values in cells between control and NGA. Two-sided Student’s t-test. *P<0.05, **<0.01, ***P<0.005, and ****P<0.001.

Notch signaling activation causes proteolytic cleavage of a Notch receptor intracellular domain. The cleaved intracellular domain translocates into the nucleus, regulating the expression of Notch-specific genes that modulate lineage-specific TF expression to instruct cell identity (5). Recent reports demonstrated that NOTCH family TFs induces ADRN-MES switching(5). Our results demonstrated that NB cells physically conforming to the ECM topography on NGA showed higher nuclear (intracellular) levels of Notch1 relative to flat controls, implying the switch towards MES (Fig. 2d, e and Supplementary Fig. 3b). These results indicate that NGA suppresses the expression of ADRN molecular signatures while increasing MES signatures to induce an identity switching towards MES.

### Mechano-biological regulation of epigenetic state decommissions ADRN CRC TF genes

We next asked if ECM topography can remodel epigenetic landscapes to drive ADRN-MES switching. Several analyses highlighted the master TFs, e.g., PHOX2B, GATA3, HAND2, and ISL1, binding to their respective super-enhancers and forming a meta-stable network of the ADRN CRC, although MES CRC TFs are less well-defined(26). Since the repression of ADRN CRCs is an essential early-phase step during ADRN-MES transition(5, 26), we reasoned to examine the collapse of the ADRN CRC by ECM topography.

Mechanical stimuli from ECM have been shown to influence chromatin conformation and gene transcription (27, 28). Thus, we investigated chromatin landscapes to probe promoters and enhancers underlying either the ADRN or MES CRC and whether they are remodeled in response to ECM topography. For this investigation, we conducted ATAC-seq (the assay for transposase-accessible chromatin with sequencing), a sequencing technique used for determining genome-wide chromatin accessibility. This accessibility indicates open and active states of chromatin, allowing for gene transcription, as opposed to closed and condensed chromatin, which is typically associated with gene silencing (29). ATAC-seq analysis of transcription start site (TSS) enrichment peaks revealed that chromatin around the TSS of ADRN signature genes was less open, and thus less accessible for transcription, for NB cells plated on NGA relative to the flat surface. The more open chromatin around the TSSs for cells on the control surface suggests that the corresponding ADRN genes are more poised for expression. (Fig. 3a,b).

**Figure 3:**
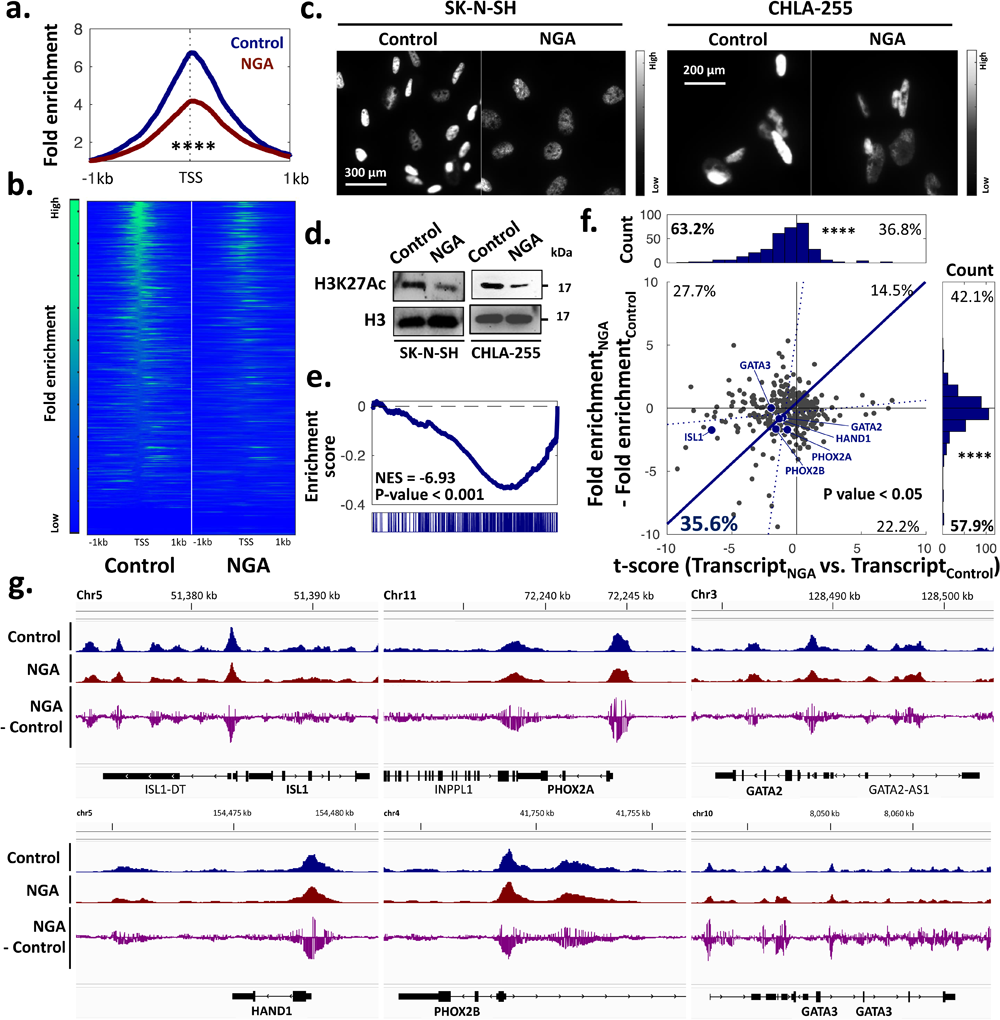
Epigenetic modulation by ECM topography decommissions ADRN CRC TF genes. Profile (**a**) and heatmap (**b**) of signal distribution around ATAC-seq open chromatin peaks around TSS regions (±1 kb) of ADRN gene signatures in SK-N-SH on control flat and NGA substrates. Higher fold enrichment, indicated by greenish than bluish, means a higher tendency to have open chromatin around TSS regions and proximal promoters in ADRN gene signatures. IF (**c**) and WB (**d**) of NB cells on control and NGA substrates with H3K27Ac antibody. **(e).** GSEA showing negative fold enrichment of open chromatin peaks in ADRN gene signatures on NGA relative to control flat substrates. This negative enrichment (or depletion) indicates SK-N-SH cells show higher open chromatin in ADRN -related genomic regions on the flat surfaces than on NGA. (**f).** Differences in the fold enrichment of ADRN signatures on control flat substrates from on those of NGA are correlated with the t-score between ADRN signature transcriptome of cells on control and NGA substrates. P-values are calculated by the Pearson correlation coefficient. A blue solid line is a linear regression. Histograms are the fold enrichment distribution of ADRN signatures (right) and the t-scores (top). Their negative values mean that the ADRN signature exhibits higher open chromatin (57.9% vs. 42.1%) and higher transcription (63.2% vs. 35.6%) on control flat than NGA substrates. Statistical difference between values and 0 and one-sample Student’s t-test. P**** < 0.001. (**g).** Open chromatin regions around ADRN CRC TF genomic regions, ISL1, PHOX2B, GATA2, HAND1, PHOX2B, and GATA3, in SK-N-SH on NGA (red) and flat (blue) surface analyzed by ATAC seq. Difference of fold enrichment in open chromatin peaks of cells between flat surfaces and NGA (purple). A negative difference indicates a lower tendency of open chromatin in the cells on NGA vs. flat surface.

We also noted a global reduction of acetylated lysine 27 residue on the H3 histone (H3K27ac) on NGA compared to on the flat substrate (Fig. 3c,d). H3K27ac indicates a relaxation of the chromatin structure. The relaxation makes the underlying DNA more accessible to transcription factors and other regulatory proteins, promoting gene transcription(29). At the same time, we noted a global increase in the repressive mark, methylated lysin 27 residue on the H3 histone (H3K27me3) associated with condensed/closed heterochromatin of cells on NGA relative to control. (Supplementary Figure. 4c). H3K27me3 is usually found at promoters of genes that are developmentally regulated and need to be kept in a silenced state during specific stages of development.

Interestingly, we found no statistically significant differences in the TSS enrichment peaks around MES signature genes, suggesting equal chromatin accessibility of MES genes in both the NGA and flat control conditions (Supplementary Figure 4a,4b). Thus, we suggest that global changes in H3K27Ac and H3K27me3 regulated by ECM topography reflect epigenetic modification in ADRN signatures specifically, not MES signatures. This finding may be consistent with a previous report suggesting that EZH2 inhibitor, suppressing H3K27me3 globally, promotes the ADRN subtype(30). Our data was corroborated further by GSEA looking at ADRN genes associated with differentially accessible peaks, showing preferential enrichment in open chromatin peaks across ADRN genes for NB cells on flat control versus NGA substrate (Fig. 3e).

**Figure 4:**
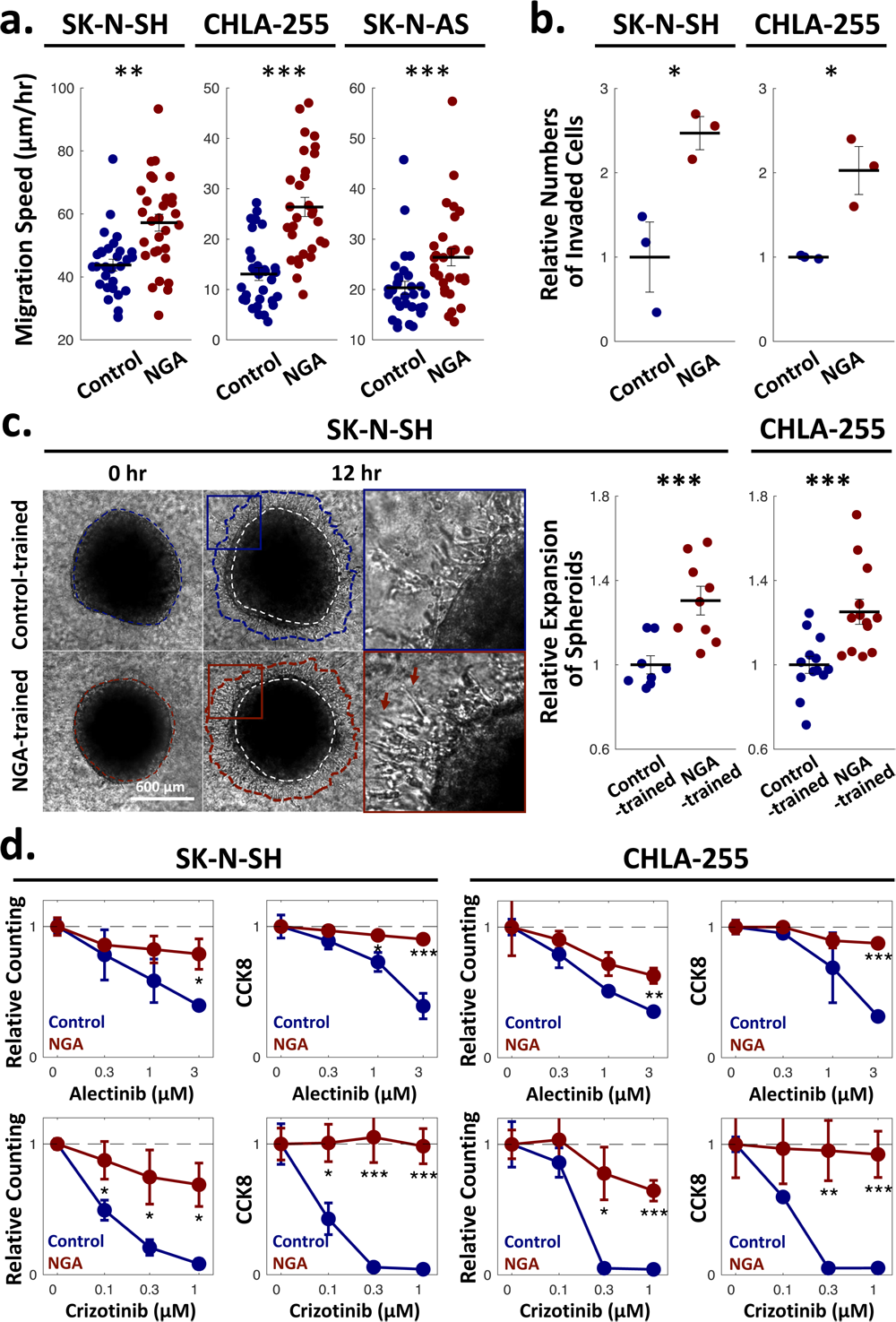
NB cells on NGA demonstrate enhanced MES phenotypical features. **(a).** On NGA, NB cells exhibit faster cell migration than on control flat substrates. **(b).** Trans-well assays with NGA-and control-trained cells, i.e., cells cultured on NGA and control substrate for 3 days, showing higher invasion of NGA-trained cells than control-trained cells. **(c).** Spheroids with NGA-trained cells exhibit more aggressive expansion than those with control-trained cells. Dashed lines (white) indicate the spheroid boundaries at the initial time point. Solid lines correspond to the spheroid boundaries after 12 hours of expansion. Relative expansion of spheroids, i.e., the ratio of the spheroid size at the initial time point to that after 12 hour-expansion. **(d).** Chemoresistance against ALK inhibitors, Alectinib and Crizotinib, of cells on control and NGA substrates. Dose-dependent survival after 2-day drug treatment was assessed by cell counting and CCK8 assay (n=3). Error bars are S.E.M. Statistical difference of cell survivals between control and NGA at each corresponding drug dose. In Panel **(a),(b),** and **(c)**, each dot indicates the value of a corresponding cell. Lines mean averages of corresponding groups. Error bars are S.E. Statistical difference of values in cells between control and NGA. Two-sided Student’s t-test. *P<0.05, P**<0.01, and ***P<0.005.

Furthermore, to examine whether the changes in chromatin accessibility across ADRN genes correlate with changes in gene expression, we plotted the differential enrichment scores from ATAC-seq data and t-scores from RNA-seq data between NGA and flat conditions. We discovered a positive clustering correlation of ADRN genes enriched in accessible chromatin peaks with transcriptional mRNA output, especially for those ADRN genes making up the ADRN CRC, namely PHOX2B, GATA3, HAND2, and ISL1 (Fig. 3f, Supplementary Table 3).

We next examined whether ECM topography decommissions promoters, enhancers, and/or other DNA regulatory elements of the ADRN CRC. We looked at the open chromatin peaks in genes serving as master TFs comprising the ADRN CRC. We noted the reduction in chromatin accessibility across ISL1, PHOX2A, GATA2, HAND1, PHOX2B, and GATA3 for cells on NGA compared to flat control. There was a general trend of promoter decommissioning for ISL1, PHOX2A, and GATA2. For HAND1, PHOX2B, and GATA3, we noted that both promoters and distal DNA regulatory elements (likely enhancers) were decommissioned in response to ECM topography (Fig. 3g.).

Collectively, this data suggests that ECM topography triggers ADRN-MES reprogramming by first epigenetically silencing the ADRN CRC through remodeling the chromatin landscape and decommissioning the respective promoters and/or enhancers of ADRN genes. However, no such epigenetic regulation was noted for MES, suggesting a transcriptional regulatory layer needed for the full induction of MES-like state.

### Switching toward MES by ECM topography is accompanied by invasiveness and drug resistance

We next investigated if the ADRN-MES reprogramming on NGA induces cells to have functional MES features, worsening tumor outcomes. Time-lapse live cell imaging revealed that NB cells on NGA show higher migration speeds than on control surfaces (Fig. 4a). Furthermore, we trained NB cells for 3 days on NGA or control surfaces and then performed a transwell invasion assay. We replated the trained cells on the permeable membrane and counted the number of invaded cells through the membrane. NGA-trained cells exhibited higher invasiveness compared to cells trained on the control substrate (Fig. 4b). Additionaly, we generated tumor spheroids with NB cells trained on NGA or control surfaces for 3 days via the hanging drop culture method and embedded them in collagen type I gels. Spheroids of NGA-trained cells exhibited aggressive expansion with spindle-like invasive cells at their margin versus spheroids of flat-trained cells (Fig. 4c and Supplementary Fig. 5).

We further found NB cells on NGA showed higher resistance to ALK inhibitors, alectinib and crizotinib, compared to cells on flat controls (Fig. 4d and Supplementary Fig. 6). This result is consistent with previous reports suggesting the MES cells in NB show resistance to ALK inhibition (6) even if they lack activating mutations in the ALK oncogene. Additionally, doxorubicin-resistant NB cells showed higher expression of MES signatures and lower expression of ADRN signatures. These results suggest that MES identity confers cells to the resistance to the conventional chemotherapeutic drug, e.g., doxorubicin, compared to control (Supplementary Fig. 6a, Supplementary Table 4). Interestingly, NB cells on NGA showed higher resistance to doxorubicin (Supplementary Fig. 7). This finding also aligns with reports suggesting that MES cells resist standard chemotherapeutic agents more than ADRN cells(1). These findings support that ECM topography causes cells to have MES phenotypes related to inferior NB outcomes.

### Rho-kinase is required for ADRN-MES transcriptional reprogramming by ECM topography

We subsequently elucidated how NB cells translate the cues from ECM topography to trigger ADRN-MES reprogramming. GSEA of NB cells plated on NGA showed statistically significant enrichment in gene sets relating to cell-ECM interactions, e.g., ECM-receptor interaction, compared to cells on flat substrates (Fig. 5a, Supplementary Table 5). When cells attach to the ECM, receptors on the cell membrane, such as integrins, cluster together. The clustering recruits other proteins, forming structures called focal adhesions. These focal adhesions serve as signaling adaptors that relay mechanical stimuli to downstream oncogenic signaling cascades. When cells encounter variations in ECM topography, such as grooves, ridges, or other patterns, the mechanical forces generated by these interactions stimulate Rho-kinase (ROCK) signaling(31, 32). ROCK triggers focal adhesion protein recruitment and stimulates their maturation. The maturation of focal adhesions reinforces cell-ECM signaling, bolstering their linkage to cytoskeleton, and promoting intracellular stress fiber formation. Interestingly, GSEA results showed higher expression of ECM-receptor interaction in MES over ADRN cells (Fig 5b). Thus, we hypothesize that ECM-receptor-mediated signaling drives the switch toward MES via ROCK signaling.

**Figure 5:**
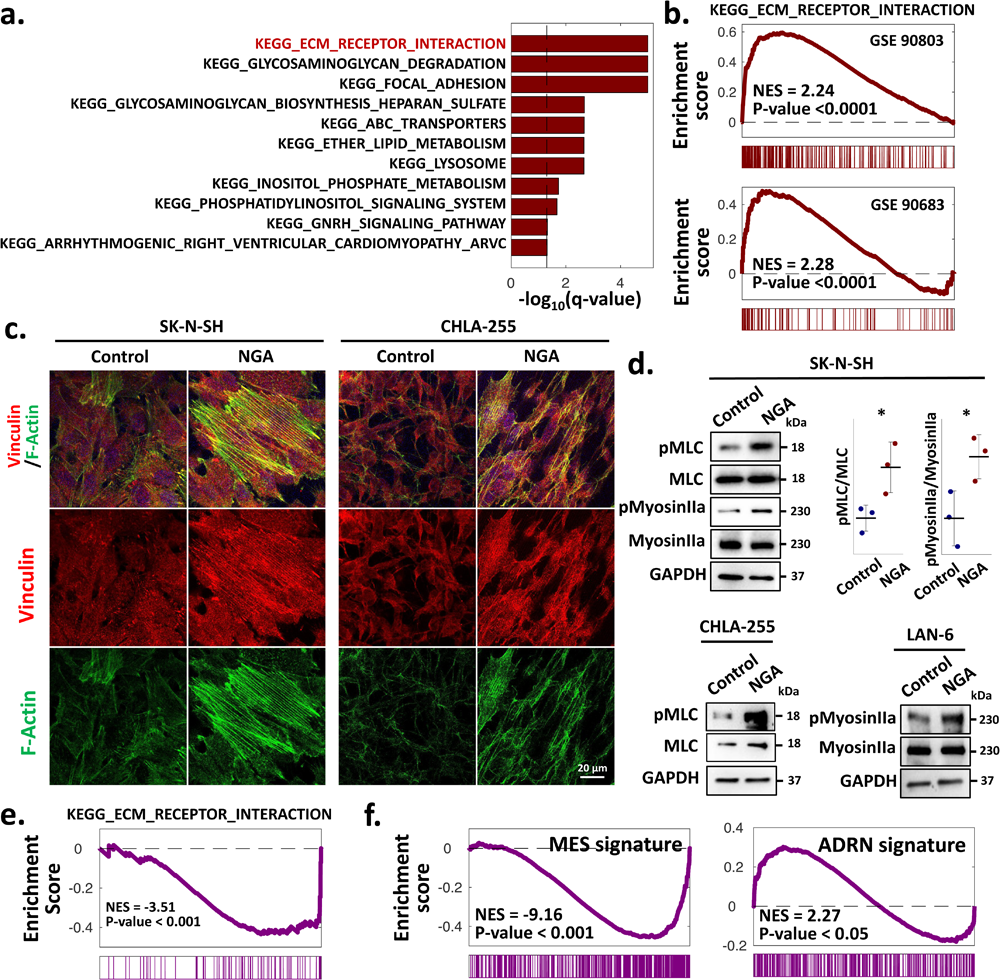
Enhanced cell-ECM interactions and ROCK activity on NGA may modulate reprogramming from ADRN to MES. **(a).** Highly enriched gene sets related to cell-ECM interactions in SK-N-SH on NGA compared to control substrates, analyzed by GSEA. Reference gene sets are KEGG. **(b).** MES exhibits positive enhancement of cell-ECM interaction-related gene sets (KEGG ECM_Receptor_Interaction), compared to ADRN. (**c).** IF of NB cells on control and NGA substrates with vinculin antibody (focal adhesion protein) and phalloidin (F-actin). **(d).** Higher ROCK activity, indicative of enhanced cell-ECM interaction in NB cells on NGA than on control substrates. Higher phosphorylation of myosin light chain (MLC) and myosin IIa implies higher actomyosin contractility. Each dot indicates the ratio of phosphorylated proteins to total proteins, indicating the phosphorylation degree of a corresponding cell. **(e).** Y27632, Rho kinase inhibitor treatment, suppressed gene expressions related to cell-ECM interaction in SK-N-SH cells on NGA. GSEA with differentially expressed genes by Y27632 (3μM, 3 hours). **(f).** Depleted MES signature expression and enhanced ADRN signature expression by Y27632 treatment (3μM) on NGA analyzed by GSEA.

To clarify if ECM topography stimulates ROCK in NB cells, we performed IF with an antibody against vinculin - a force-bearing focal adhesion protein, and phalloidin that stains fibrous-actin (F-actin) cytoskeleton, i.e., stress fiber (Fig. 5c and Supplementary Figure 8a). NB cells conforming to NGA showed stress fiber formation typified by pronounced F-actin stress fibers oriented along the grooved features of NGA together with the enhanced assembly of vinculin-enriched focal adhesions. Furthermore, we confirmed higher phosphorylation of myosin light chain (pMLC), a ROCK substrate, and myosin IIa (pMyosinIIa), a downstream molecule of MLC, of NB cells cultured on NGA versus the flat control substrate (Fig. 5d and Supplementary Fig. 8b). These results clarify that ECM topography stimulates ROCK and lead us to investigate further if ROCK mediates ADRN-MES transition.

Thus, we explored if ROCK inhibition prevents ADRN-MES transition. We first confirmed that the treatment of NB cells on NGA with the Rho kinase inhibitor (ROCKi), Y27632 (3μΜ for 3hr), reversed the previously observed enrichment of gene sets relating to cell-ECM interactions (Fig 5e) via RNA seq followed by GSEA. In addition, GSEA showed that Y27632-treated NB cells failed to undergo transcriptional ADRN-MES reprogramming on NGA and retained ADRN-specific signatures (Fig. 5f and Supplementary Fig. 8c, Supplementary Table 6). Overall, this data suggests that NB cells conforming to ECM topography display enhanced cell-ECM interactions via ROCK signaling stimulation. ROCK signaling may serve as a signaling hub of topography sensing and mechano-transduction that is necessary for driving ADRN-MES reprogramming.

### ROCK inhibition reverses ADRN-MES reprogramming by ECM topography

We next asked whether ROCK inhibition can abolish ADRN-MES transition on NGA. NB cells plated on NGA and subsequently treated with ROCKi, Y27632, failed to undergo the molecular changes associated with MES switching. This was evident by a decrease in expression of MES markers, PRRX1 and NOTHC1, and an increase in expression of ADRN markers, GATA3 and PHOX2B. We also observed a global increase in the expression levels of H3K27ac (Fig 6a-d) by ROCKi treatment. Also, the treatment suppressed cell migration speed of NB cells on NGA and spheroid expansion of NGA-trained cells through 3D collagen matrices (Fig 6e,f and Supplementary Fig. 9). Similarly, ROCKi reversed the drug resistant phenotype on NGA and partially re-sensitized NB cells to the ALK inhibitors (Fig. 6g). These results suggest that ROCK inhibition, suppressing cell-ECM interaction, prevents ADRN-MES phenotypic switching cells on NGA by blocking mechanotransduction of topographical ECM cues.

**Figure 6:**
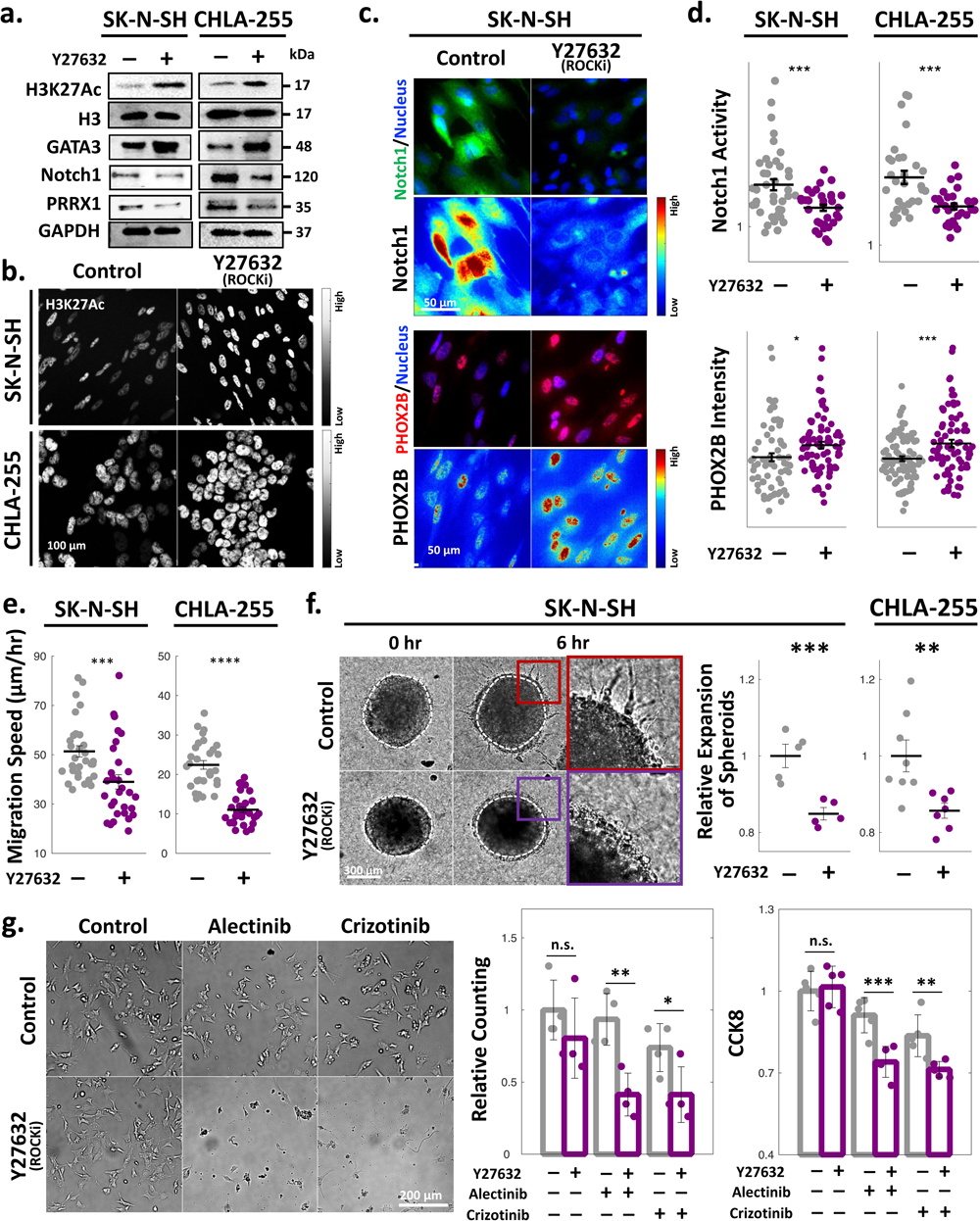
ROCK inhibition restores ADRN features on NGA. **(a).** WB of NB cells on NGA with and without Y27632 treatment (3μM), suggesting that ROCK inhibition stimulates ADRN CRC via epigenetic modulation indicated by increased H3K27Ac and GATA3 and prevents reprograming toward MES shown as reduced intracellular Notch1 and PRRX1. **(b).** IF of Y-27632 treated NB cells on NGA exhibits higher H3K27Ac than non-treated cells. **(c).** IF of Y-27632 treated NB cells on NGA with antibodies of Notch 1 (top) and PHOX2B (bottom). Pseudo-color corresponds to IF intensity. **(d)** Quantification of Notch1 activity (quantified by the ratio of nuclear Notch1 IF intensity to cytoplasmic Notch1 IF intensity) and PHOX2B IF intensity of cells on NGA with and without Y27632 treatment. Decreased cell migration speeds (**e**) and spheroid expansion for 6 hours (**f**) by Rho kinase inhibition. White dashed lines indicate spheroid boundaries at the initial time point. Relative expansion of spheroids, i.e., the ratio of the spheroid size at the initial time point to that on 6 hours after Y27632 treatment. **(g).** Cell survival of NGA-trained cells with Y27632 against ALK inhibitors, analyzed by cell counting and CCK8. We cultured SK-N-SH cells on NGA for 2 days and replated the trained cells. Then, we treated Y27632 along with ALK inhibitors, Alectinib and Crizotinib. In Panel **(d),** (**e),** and **(f)**, each dot indicates the value of a corresponding cell. Lines mean averages of corresponding groups. Error bars are S.E. Statistical difference of values in cells between control and NGA. Two-sided Student’s t-test. *P<0.05, P**<0.01, and ***P<0.005.

## Discussion

The relative paucity of recurrent mutations in the NB guides us to explore TME as extrinsic cues that modulates NB heterogeneity and clinical outcome(26). Our findings reveal that high-risk and relapsed NB show stromal enrichment of topographically aligned collagen-III fibrils, suggesting its potential effect on NB progression. Using an *in vitro* aligned ECM model, we demonstrate that the distinct topographical ECM features stimulate ADRN-MES transition by ROCK signaling.

Specifically, we introduce that mechano-biological reprogramming by ECM topography involves rewiring of the epigenetic state of NB cells towards MES. To our knowledge, this is the first time to demonstrate that environmental physical cues from ECM can epigenetically disrupt the metastable arrangement of the ADRN CRC by altering chromatin accessibility and decommissioning ADRN master TF genes. However, we didn’t observe changes in the occupancy of MES signature. Instead, we discovered that a transcriptional upregulation of PRRX1 and higher activation of Notch signaling by ECM topography that can trigger MES reprogramming after the ADRN CRC is epigenetically silenced.

Rho-associated kinase has attracted greater interest as a therapeutic NB target because 39% of high-risk NB patients had mutations in genes regulating Rho/Rac signaling (33). The mutations are associated with activation of the downstream ROCK and higher ROCK2 expression correlates with poor prognosis(34). Selective inhibition of the ROCK2 isoform with the small molecule HA1077 has been shown to promote MYCN protein degradation, induce neuroblastoma cell differentiation, and repress neuroblastoma growth *in vitro* and *in vivo*. Our results also support a therapeutic effect of ROCK inhibition. Y27632, exerting equal inhibitory action against both ROCK1 and ROCK2 isoforms, suppresses mechano-sensing of the biophysical environment and ADRN-MES transition. Thus, further investigation will be needed to explore how our findings about ROCK inhibition apply to an *in vivo* setting.

Overall, our findings suggest that the TME factor, i.e., ECM topography, aggravates NB outcomes by inducing ADRN-MES reprogramming. The mechano-biological reprogramming involved rewiring the epigenetic and transcriptional landscape accompanied by resistance to targeted therapy (ALKi) and increased invasiveness. Our study opens new avenues for exploration in this pediatric malignancy highlighting the role of the ECM and mechanotransduction as key regulators of the cellular heterogeneity and phenotypic plasticity underlying tumor aggressiveness in high-risk NB.

## Methods

### Cell lines, antibodies and drugs

NB cell lines SK-N-SH and SK-N-AS were purchased from ATCC. CHLA-255 and LAN-6 were a kind gift from Dr. Shahab Asgharzadeh (Children’s Hospital Los Angeles, CA). Cell lines were cultured in high glucose DMEM supplemented with 20% FBS and 1% penicillin/streptomycin. We authenticated cell lines via STR profiling (CHLA Stem Cell core). Drugs and inhibitors were as followed: Y27632 ROCK inhibitor (SCM075 from Sigma Aldrich), Alectinib (S2762 from SelleckChem), Crizotinib (PZ0191 from Sigma-Aldrich), Doxorubicin hydrochloride (D1515 from Sigma-Aldrich). Antibody list is as followed:

| Antibody List | Species | Vendor | Catalogue Number | Dilution | Assay |
| --- | --- | --- | --- | --- | --- |
| Anti-Collagen III antibody [FH-7A] | Mouse monoclonal | abcam | ab6310 | Dilution 1:300 | Immunohistochemistry |
| Anti-PRRX1 antibody | Mouse monoclonal | Sigma-Aldrich | SAB1412737 | 2.5 ug/mL | Western Blotting |
| Recombinant Anti-GATA3 antibody [EPR16651] | Rabbit monoclonal | abcam | ab199428 | Dilution 1:1000 | Western Blotting |
| Recombinant Anti-PHOX2B antibody [EPR14423] | Rabbit monoclonal | abcam | ab183741 | Dilution 1:100 | IF |
| Anti-Vimentin antibody [RV202] | Mouse monoclonal | abcam | ab8978 | 1 ug/mL | IF |
| Anti-NOTCH1 (D1E11) | Rabbit monoclonal | CST | #3608 | Dilution 1:1000 or 1:100 | Western Blotting/IF |
| Acetyl-Histone H3 (Lys27) (D5E4) | Rabbit monoclonal | CST | #8173 | Dilution 1:1000 or 1:100 | Western Blotting |
| Tri-Methyl-Histone H3 (Lys27) (C36B11) | Rabbit monoclonal | CST | #9733 | Dilution 1:1000 | Western Blotting |
| Anti-Histone H3 antibody | Rabbit polyclonal | abcam | ab1791 | Dilution 1:1000 | Western Blotting |
| Anti-Vinculin antibody | Rabbit polyclonal | Sigma-Aldrich | V4139 | Dilution 1:250 | IF |
| Phospho-Myosin IIa (Ser1943) Antibody | Rabbit | CST | #5026 | Dilution 1:1000 | Western Blotting |
| Myosin IIa Antibody | Rabbit | CST | #3403 | Dilution 1:1000 | Western Blotting |
| Myosin Light Chain 2 Antibody | Rabbit | CST | #3672 | Dilution 1:1000 | Western Blotting |
| Phospho-Myosin Light Chain 2 (Thr18/Ser19) Antibody | Rabbit | CST | #3674 | Dilution 1:1000 | Western Blotting |
| GAPDH (D4C6R) | Mouse monoclonal | CST | #97166 | Dilution 1:1000 | Western Blotting |

### Nanogroove array (NGA) biomaterial fabrication

The nanostructured substrata (width: 1000 nm, spacing: 1000 nm, and height: 1000 nm) were fabricated by capillary-force lithography with UV-curable polyurethane acrylate (PUA, Minuta Technology, South Korea) polymer having the rigidity value of 0.1 GPa, unless indicated otherwise. To fabricate a flexible polymer mold, a precursor of UV-curable polymer was drop-dispensed onto the prefabricated silicon master with large areas of 25 × 25 mm2. Then a polyethylene terephthalate (PET) (SKC Inc., South Korea) film with thickness of 50 μm was brought into conformal contact onto the polymer resin. After removing air bubbles with a roller, UV (λ = 250–400 nm) was irradiated for few tens of seconds for the cross-linking. The flexible and transparent mold with PET support was obtained after peeling off from the master, leaving behind regularly spaced nanogrooves, i.e., NGA. For the polymer nanogroove patterns on the slide glass-sized coverslips (50 × 24 mm, Thermo Fisher), the same replication process was performed onto a cleaned coverslip (circular, 12 mm diameter or rectangular 24 x 50 mm), using the replicated PUA pattern as a mold. The flat surfaces were generated as a control experimental set with the same procedure except no master. For cell-culture studies, both NGA and flat control substrates were pre-coated with collagen type III (Advanced Biomatrix) at a concentration of 50 ug/mL in a 0.01M HCL solution for 1 hour at room temperature.

### Gene set enrichment assay with the transcriptome of MES cells compared to ADRN cells

Transcriptome data of MES and ADRN NB cells are from GSE90683 and GSE90803. Differentially expressed genes between MES and ADRN cells, which adjust p-value <0.05, are identified by two-paired Student’s t-test followed by measurement of false discovery rate, which calculated adjust p-values. We performed GSEA with the identified differentially expressed genes via GSEA software (https://www.gsea-msigdb.org/gsea/index.jsp) or a customized Matlab (MathWorks, Natick, MA) code based on Subaramanian *et al.*(35). Briefly, we calculated tscores of each gene expressed in MES and ADRN cells. Then, we applied the ranked genes based on t-scores for GSEA. Ranked genes in GSE90683 and GSE90803 and their GSEA reports are in Supplementary table 1.

### Quantification of ECM alignment

10 formalin fixed paraffin embedded (FFPE) tumor tissues from patients diagnosed with neuroblastoma were evaluated by immunohistochemistry (IHC) for Collagen III expression (clone FH-7A, Abcam, 1:300 dilution). The samples included diagnostic tissue from three low risk, stage 1 tumors; two high risk, stage 4 tumors with MYCN amplification; two high risk, stage 4 tumors without MYCN amplification; as well as 3 tumor samples collected at relapse. Collagen fiber alignment was quantified with image segmentation analysis from FFPE tissue sections. We segmented collagen type III fibers in IHC images with imbinarize function of Matlab (MathWorks, Natick, MA) and assessed the orientation angles of the segmented fibers.

### Scanning Electron Microscopy (SEM)

For scanning electron microscopy, cultured cells on NGA were washed with phosphate-buffered saline (PBS, pH 7.4, Gibco Invitrogen) and fixed in 3% glutaraldehyde (Sigma-Aldrich) in PBS for 1 hr. After fixation, we rinsed samples in 0.1 M sodium cacodylate for thirty minutes at 4°C and then post-fixed in 2% osmium tetroxide for 1 hr at the same temperature. After a brief D-H2O rinse, samples were en-bloc stained in 2% aqueous uranyl acetate (0.22 μm filtered) for 1 hr at room temperature in the dark. Following a graded ethanol dehydration cells were critical point dried with liquid CO2, mounted onto SEM stubs with double stick carbon tape, and sputter coated with 10 nm gold palladium. Samples were viewed and photographed with a LEO FESEM 1530 operating at 1 kV.

### RNA sequencing

RNA was extracted from cells cultured on control or NGA surfaces using the Qiagen RNeasy kit. RNA library preparations and sequencing reactions were conducted at Azenta Life Sciences (South Plainfield, NJ, USA). RNA samples were quantified using Qubit 2.0 Fluorometer (Life Technologies, Carlsbad, CA, USA) and RNA integrity was checked using Agilent TapeStation 4200 (Agilent Technologies, Palo Alto, CA, USA). The RNA sequencing libraries were prepared using the NEBNext Ultra II RNA Library Prep Kit for Illumina using manufacturer’s instructions (New England Biolabs, Ipswich, MA, USA). Briefly, mRNAs were initially enriched with Oligod(T) beads. Enriched mRNAs were fragmented for 15 minutes at 94°C. First strand and second strand cDNA were subsequently synthesized. cDNA fragments were end repaired and adenylated at 3’ends, and universal adapters were ligated to cDNA fragments, followed by index addition and library enrichment by PCR with limited cycles. The sequencing libraries were validated on the Agilent TapeStation (Agilent Technologies, Palo Alto, CA, USA), and quantified by using Qubit 2.0 Fluorometer (ThermoFisher Scientific, Waltham, MA, USA) as well as by quantitative PCR (KAPA Biosystems, Wilmington, MA, USA). The sequencing libraries were clustered on one flowcell lane. After clustering, the flowcell was loaded on the Illumina HiSeq instrument (4000 or equivalent) according to manufacturer’s instructions. The samples were sequenced using a 2×150bp Paired End (PE) configuration. Image analysis and base calling were conducted by the Control software.

Raw sequence data (.bcl files) generated from the sequencer were converted into fastq files and de-multiplexed using Illumina’s bcl2fastq 2.17 software. One mismatch was allowed for index sequence identification. After investigating the quality of the raw data, sequence reads were trimmed to remove possible adapter sequences and nucleotides with poor quality using Trimmomatic v.0.36. The trimmed reads were mapped to the reference genome available on ENSEMBL using the STAR aligner v.2.5.2b. The STAR aligner uses a splice aligner that detects splice junctions and incorporates them to help align the entire read sequences. BAM files were generated as a result of this step. Unique gene hit counts were calculated by using feature Counts from the Subread package v.1.5.2. Only unique reads that fell within exon regions were counted. The counted numbers of SK-N-SH cells on the flat control and NGA are in Supplementary table 2 and those of SK-N-SH cells with ROCKi are in Supplementary table 3. After extraction of gene hit counts, the gene hit counts table was used for downstream differential expression analysis. We identified differentially expressed genes between cells on NGA and on the flat control surface (Supplementary table 4) and between ROCKi treatment and without the treatment and perform GSEA with the identified genes as above. Principial component analysis was performed with PCA function of Matlab (MathWorks, Natick, MA). Also, we calculated t-scores of genes expressed differently between NGA and the flat control surfaces and between control and ROCKi treatment and performed GSEA with the ranked genes based on the t-scores. MES and ADRN signatures are from Groningen *et al* (3).

Transcriptomes of doxorubicin resistant NBL cells are acquired from GSE47670 (REF). We ranked genes based on difference in gene expression between control and doxorubicin resistant NBL cells using R2: Genomic Analysis and Visualization Platform (https://hgserver1.amc.nl/) and performed GSEA (Supplementary table 5).

### ATAC sequencing

Cells were trypsinized from either flat or NGA surfaces and centrifuged at 600g for 5 min at 4C. After removing the supernatant, cells were resuspended in 500 uL growth media +10% DMSO and solutions was transferred to a 1.7mL microcentrifuge tube on ice. Tubes containing the suspended cells were added to Mr. Frosty and stored at −80C after 24hr for cryopreservation.

ATAC-seq library preparation and sequencing reactions were conducted at Azenta Life Sciences (South Plainfield, NJ, USA). Live cell samples were thawed, washed, and treated with DNAse I (Life Tech, Cat. #EN0521) to remove genomic DNA contamination. Live cell samples were quantified and assessed for viability using a Countess Automated Cell Counter (Thermo Fisher Scientific, Waltham, MA, USA). After cell lysis and cytosol removal, nuclei were treated with Tn5 enzyme (Illumina, Cat. #20034197) for 30 minutes at 37°C and purified with Minelute PCR Purification Kit (Qiagen, Cat. #28004) to produce tagmented DNA samples. Tagmented DNA was barcoded with Nextera Index Kit v2 (Illumina, Cat. #FC-131-2001) and amplified via PCR prior to a SPRI Bead cleanup to yield purified DNA libraries.

The sequencing libraries were clustered on a single lane of a flowcell. After clustering, the flowcell was loaded on the Illumina HiSeq instrument (4000 or equivalent) according to manufacturer’s instructions. The samples were sequenced using a 2×150bp Paired End (PE) configuration. Image analysis and base calling were conducted by the HiSeq Control Software (HCS). Raw sequence data (.bcl files) generated from Illumina HiSeq was converted into fastq files and de-multiplexed using Illumina’s bcl2fastq 2.17 software. One mismatch was allowed for index sequence identification. After investigating the quality of the raw data, sequencing adapters and lowquality bases were trimmed using Trimmomatic 0.38. Cleaned reads were then aligned to reference genome hg38 using bowtie2. Aligned reads were filtered using samtools 1.9 to keep alignments that (1) have a minimum mapping quality of 30, (2) were aligned concordantly, and (3) were the primary called alignments. PCR or optical duplicates were marked using Picard 2.18.26 and removed. Prior to peak calling, reads mapping to mitochondria (mt) were called and filtered, and reads mapping to unplaced contigs were removed.

MACS2 2.1.2 was used for peak calling to identify open chromatin regions. Valid peaks from each group or condition were then merged and peaks called in at least 66% of samples were kept for downstream analyses. For each pair-wise comparison, peaks from condition A and condition B were merged and peaks found in either condition were kept for downstream analyses. Reads falling beneath peaks were counted in all samples, and these counts were used for differential peak analyses using the R package Diffbind (Supplementary table 6). We performed GSEA with the ranked genes based on the difference in fold enrichment. MES and ADRN signatures are from Groningen *et al* (3).

### Western Blotting

NB cells were lysed from their respective substrates and collected in RIPA buffer containing protease/phosphatase inhibitors (Halt Protease/Phosphatase Inhibitor Cocktail 100x). Protein lysates were centrifuged at 14,000g for 15 min to pellet cell debris, and the supernatant was removed and quantified with BCA assay. Protein samples were run using SDS-PAGE (Mini-PROTEAN TGX Gels, 4-20%, Bio-Rad) and transferred to polyvinylidene fluoride (PVDF) membranes. Membranes were blocked in 3% BSA (Sigma) in TBST for 1 hour and then incubated with primary antibodies. Primary antibodies (10 μg per well) include PRRX1, GATA3, NOTCH1, MLC2, pMLC2 (Ser 19), Myosin-IIa, pMyosin-IIa, H3K27ac, H3K27me3, H3, GAPDH. Refer to Supplementary Table 2 for all antibody information. All antibodies were used at a 1:1,000 dilution and incubated overnight at 4C. Next, membranes were washed 3 times in TBST for 5 min each to remove unbound antibody and incubated with HRP-conjugated IgG secondary antibodies (1:2000-1:5000) for one hour at room temperature. Membranes were washed again 3 times with TBST for 5 min each to remove unbound secondary antibody. Protein bands were visualized using Clarity Western ECL substrate (Bio-Rad) using the iBright FL1500 imaging system.

### Immunofluorescence and quantification

NB cells were seeded on collagen-III coated NGA or flat substrates. Following pertinent treatment, cells were fixed with 4% paraformaldehyde in PBS for 10 min, permeabilized with 0.1% Triton X-100 (Sigma) in PBS for 10 min and blocked with 1% BSA-PBS (Sigma) for 45 min. For actin-cytoskeleton staining, samples were incubated with phalloidin-Atto 647N (Sigma) for 2 h. Primary antibodies diluted in blocking solution (1:100-1:250) were incubated for 2hr and washed 3 times with PBS to remove unbound antibody. We followed by 1 h incubation at room temperature with Alexa 488-and/or Alexa 594-labelled secondary antibodies (Invitrogen) at a concentration of 5mg/mL in blocking solution. Nuclei were counterstained with 4’,6-diamino-2-phenylindole (Prolong Gold antifade reagent DAPI; Invitrogen). The primary antibodies were vimentin, phox2b, notch1, H3K27ac. Confocal and epifluorescence images were collected using a Nikon Eclipse Ti2 inverted microscope. We analyzed the images using a customized MATLAB (The MathWorks, Natick, MA) code to read out the intensity of each pixel, indicating the amounts of corresponding proteins. Statistical analyses were performed using unpaired twosided Student’s *t*-tests. At least 20 cells of each experimental condition were assessed.

### Cell migration speed assay

We assembled the patterned rectangular coverslip onto 8-multiwell chambers (Nunc® Lab-Tek® Chamber SlideTM system) after removing pre-mounted slide glasses on their bottom. After coating the pattern with collagen-III, NB cells were replated and recorded for 10 hours, using Nikon C2+ confocal microscope system with a controllable mechanical stage allowing X-Y motion and Piezo Z-stage control, DS-Qi2 monochrome CMOS camera, as well as environmental control for live-cell imaging. Cells were manually traked in every image taken every 20 min in a time-lapse imaging for 10 h, using a customized MATLAB (The MathWorks, Natick, MA) code. The tracked time-series coordinates of cells were used to calculate cell migration speeds by averaging displacements per each time interval, 20 min. Statistical analyses were performed using unpaired two-sided Student’s *t*-tests. Every experimental condition was analyzed in duplicate and at least 40 cells were tracked in each data replicate.

### Transwell assay

Cell invasiveness was measured using transwell chambers (Millicell 24 Cell Culture Insert Plate PCF 8 μm). NB cells trained on NGA and flat substrates for 3 days were trypsinized, resuspended in serum-free DMEM media, and added to the upper chamber with the membrane (8 μm pore size) in the insert. Then, the inserts were placed in the lower chamber containing media with 20% FBS. Following incubation at 37C for 24 hr, cells that migrated to the lower chamber were fixed with 4% paraformaldehyde in PBS and stained with Hoesht dye. Stained cells were visualized with epifluorescence imaging. We counted the number of cells per 20x magnification field. Statistical analyses were performed using unpaired two-sided Student’s *t*-tests. At least 3 images of every experimental condition were analyzed.

### Hanging drop 3D tumor spheroid and collagen invasion assay

NB cells were pre-trained on collagen-III coated (50 μg/mL) NGA or flat substrate for 3 days. To generate spheroid aggregates, the hanging drop method was employed. In short, pre-trained cells were trypsinized and made into a suspension at a density of 10^6^ cells/mL. As many as 15 total 20uL drops (20,000 cells in each drop) were deposited across the lid of a 60mm petri dish. The lid with deposited drops was carefully inverted and placed on top of the PBS-filled bottom chamber of the petri dish. The hanging drops were maintained in a 37C, 5% CO2 incubator for 48 hours until spheroids were formed.

A 3mg/mL stock rat tail collagen I solution (Gibco, Life Technology) was used to make a total volume of 1 mL of pH-buffered collagen gel solution at a final density of 2mg/mL. In a sterile tube 213 μL of dH2O, 100 μL 10xDMEM, and 17 μL 1N NaOH along with 670 μL of collagen I solution was added and briefly shaken until color of solution turns from yellow to slight pink. The collagen-I solution was kept on ice to prevent polymerization.

The spheroids formed in hanging drops were then collected in growth media, centrifuged at low speed 100g for 3 min, resuspended in growth media and embedded in the rat tail collagen-I solution (Gibco, Life Technology) with a 1:3 ratio (cells: gel mixture) and deposited in wells of a glass-bottom 24 well plate. The plate was then incubated for 1 hour to let the 3D gel:cell mixture polymerize and media was then added on top. Time-lapse imaging (brightfield) of the tumor spheroids embedded in 3D collagen-I gels was conducted to monitor their expansion dynamics.

### Drug resistance assay

NB cells were seeded on collagen-III coated NGA 96 well plates (CuriBio) or flat control plates at a density of 10,000 cells per well. Drug treatment was initiated 24hr after cell seeding. Alectinib (Selleck Chemicals) was added at concentration of 0, 0.3, 1 and 3 μM in technical triplicates. Crizotinib (Sigma) was added at concentration of 0, 0.1, 0.3 and 1 μM in technical triplicates Doxorubicin (Selleck Chemicals) was added at concentration of 0, 0.1, 0.3, 1 and 3 μM in technical triplicates. Duration of drug treatment was 48hr before determining cell viability with CCK-8 assay (Sigma). The amount of the formazan dye generated by reduction of the tetrazolium salt WST-8 by the metabolic activity of dehydrogenases in cells is directly proportional the number of living cells and quantified with a microplate reader at 450 nm absorbance. Alternatively, cell viability was also determined with optical brightfield microscopy by counting number of cells across 3 different field of views per well at a 10x magnification each.

### Supplementary Tables

Supplementary Table 1: Gene Set Enrichment Analysis (GSEA) from REACTOME for GSE90803/GSE90683

Supplementary Table 2: Differential Gene Expression (DGE) from RNA-seq of SK-N-SH on Control/NGA

Supplementary Table 3: ATAC-seq with annotated peaks for SK-N-SH on Control/NGA

Supplementary Table 4: Gene Set Enrichment Analysis (GSEA) for doxorubicin-resistant cells

Supplementary Table 5: Gene Set Enrichment Analysis (GSEA) from KEGG for SK-N-SH on Control/NGA

Supplementary Table 6: Differential Gene Expression (DGE) from RNA-seq of SK-N-SH on NGA treated with Y27632 (3μM)

### Source Data

Source Data 1: Raw data for Supplementary Figure 2

Source Data 2: Raw data for Figure 2

Source Data 3: Raw data for Figure 3f

Source Data 4: Raw data for Figure 4, Supplementary Figure 5 and Supplementary Figure 7c

Source Data 5: Raw data for Figure 5d

Source Data 6: Raw data for Figure 6f, g

---

**Supplementary Figure 1.**
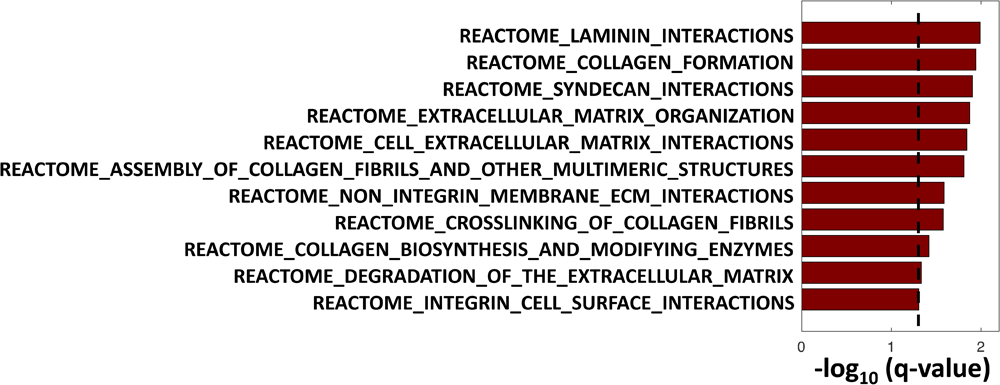
GSEA REACTOME of MES versus ADRN. GSEA analysis of transcriptomic data from the Reactome database (GSE90803) shows MES cells, compared to ADRN cells, are highly enriched in gene sets relating to pathways involved in cell-ECM interactions, ECM (collagen and laminin) formation, and organization. A value of negative log10(q-value) more than 1 was defined as significant, and a higher value indicates stronger enrichment.

**Supplementary Figure 2.**
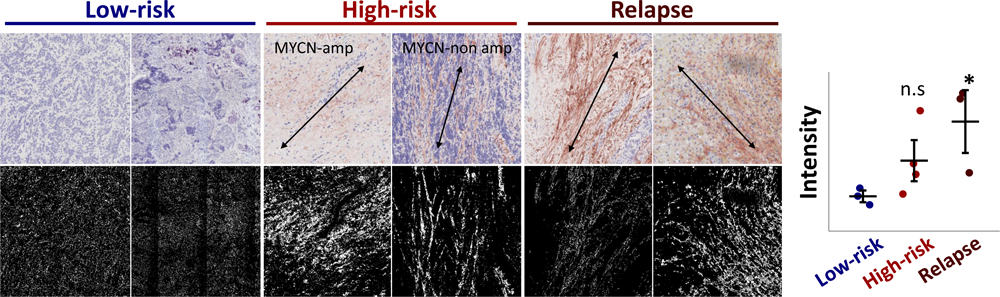
Col-III immunohistochemical staining on risk-stratified NB tissues. Immunohistochemistry with Col-III antibody (clone FH-7A, Abcam, 1:300 dilution) was performed on formalin-fixed paraffin-embedded (FFPE) tissues from patients diagnosed with NB. Upper panel showing representative staining intensity for low-risk stage 1 tumors (n=3), high-risk stage 4 tumors with MYCN amplification (n=2), high-risk stage 4 tumors without MYCN amplification (n=2) as well as tumors collected at relapse (n=3). Arrows represent the orientation angle of col-III fibrils. The lower panel highlights the topographical alignment of Col-III fibrils. Staining intensity quantification for Col-III shows a higher intensity score for high-risk and relapsed NB tissues which reached statistical significance only for relapsed disease. Two-sided Student’s t-test. *P<0.05, **<0.01, ***P<0.005, and ****P<0.001.

**Supplementary Figure 3.**
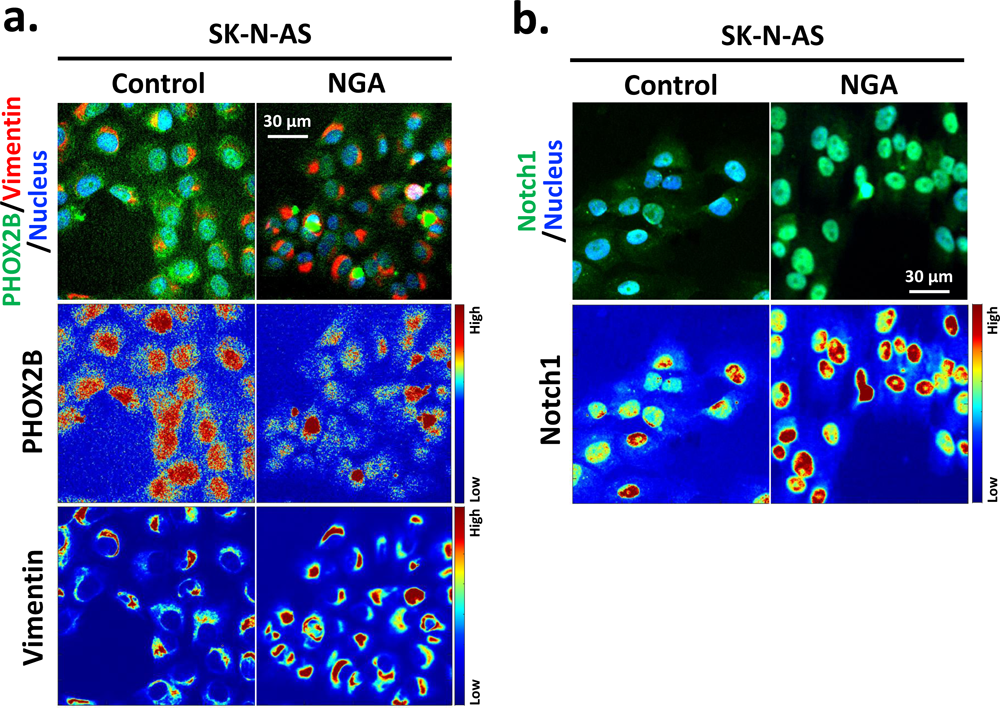
Increased MES and decreased ADRN marker expression for SK-N AS on NGA. (**a**) IF staining against PHOX2B (ADRN marker) and Vimentin (MES marker) on NGA versus control. Pseudo-color images indicate IF intensity showing lower nuclear levels of PHOX2B and higher cytoskeletal vimentin levels on NGA relative to control (**b**) IF staining against NOTCH1 (MES marker) showing increased nuclear NOTCH1 expression on NGA relative to control.

**Supplementary Figure 4.**
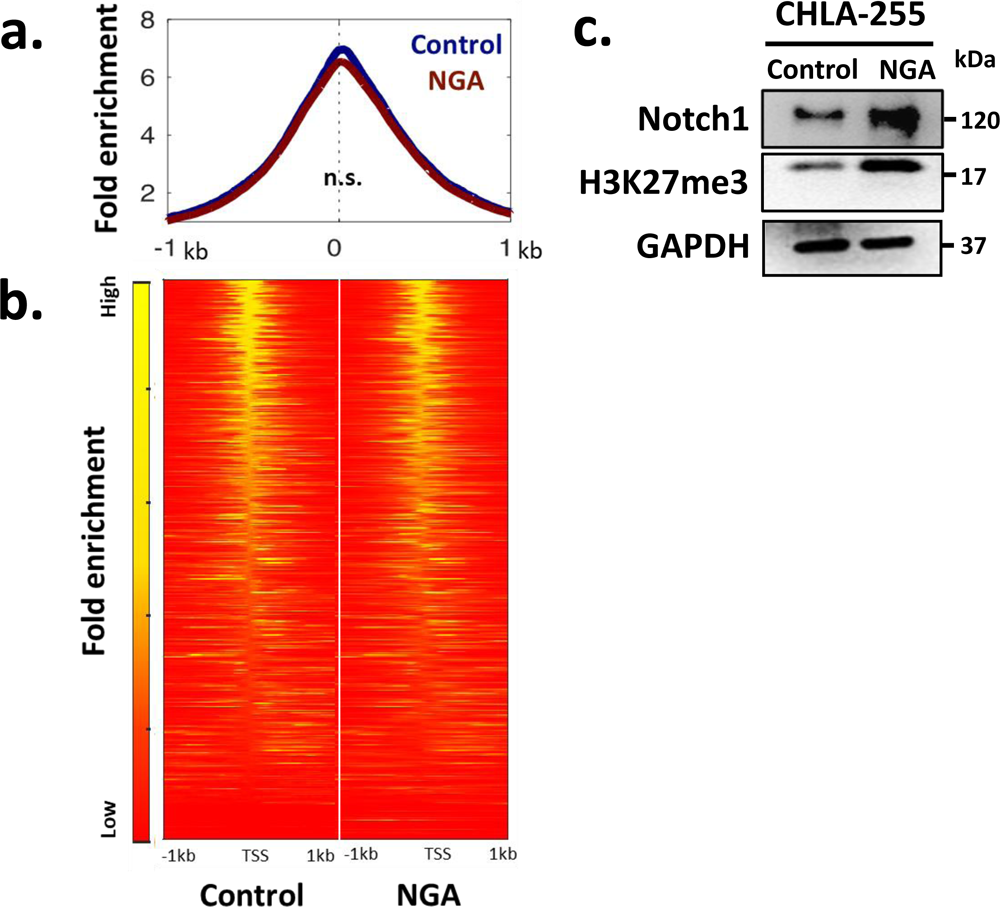
MES CRC TF genes show no mechano-biological epigenetic regulation by ECM topography. **(a)** Enrichment of open chromatin peaks from ATAC-seq data on TSS regions (±1 kb) and proximal promoters of MES CRC TF genes show no statistical enrichment for NB cells cultured on NGA relative to control. **(b)** Heat maps of signal distribution of open chromatin on TSS regions (±1 kb) of MES CRC TF genes. (**c**) Western blotting of CHLA-255 cells on NGA show an increase in the intracellular domain of Notch1 (120 kDa) which is an MES marker as well a global increase in the repressive heterochromatin mark H3K27me3 expression level relative to controls.

**Supplementary Figure 5.**
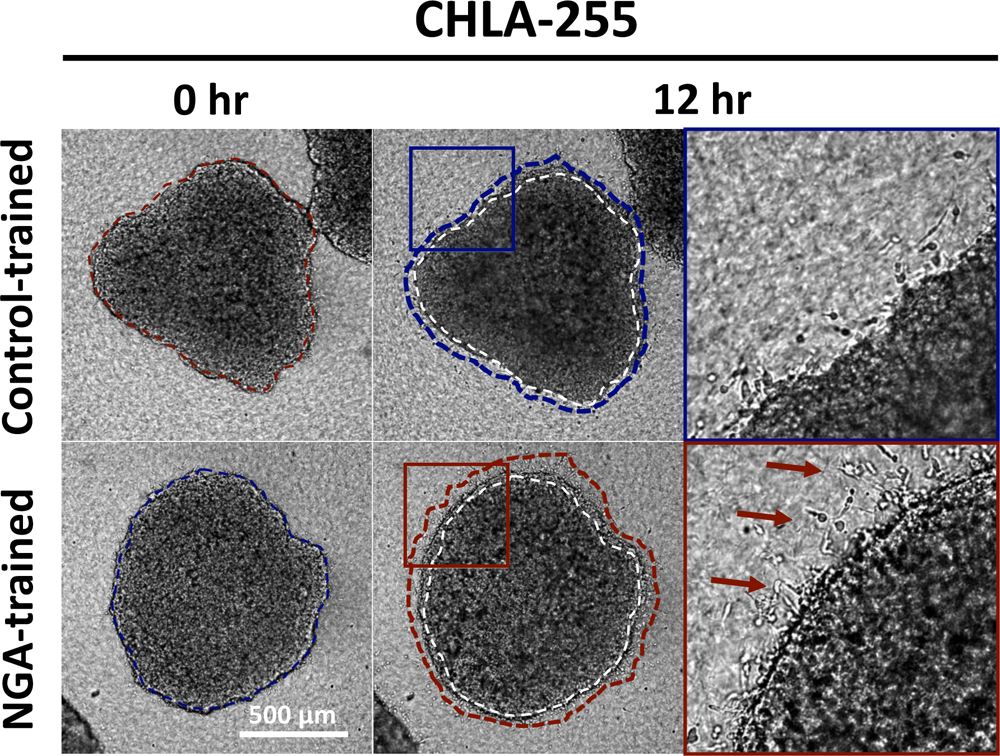
NGA-trained CHLA-255 tumor spheroids show increased invasiveness in 3D collagen-I gels. CHLA-255 cells were pre-trained on NGA and control substrates for 3 days and subsequently grown in spheroids via the hanging drop method. NGA and control-trained spheroids were embedded in 3D collagen gels and their expansion dynamics were monitored with time-lapse imaging for 12 hr. Increased sprouting and invasive spreading were noted for NGA-trained spheroids. White and red dashed lines represent the relative size of spheroids at t=0 and t=12 hr respectively. Red arrows indicate invading cells at the tumor-ECM boundary.

**Supplementary Figure 6.**
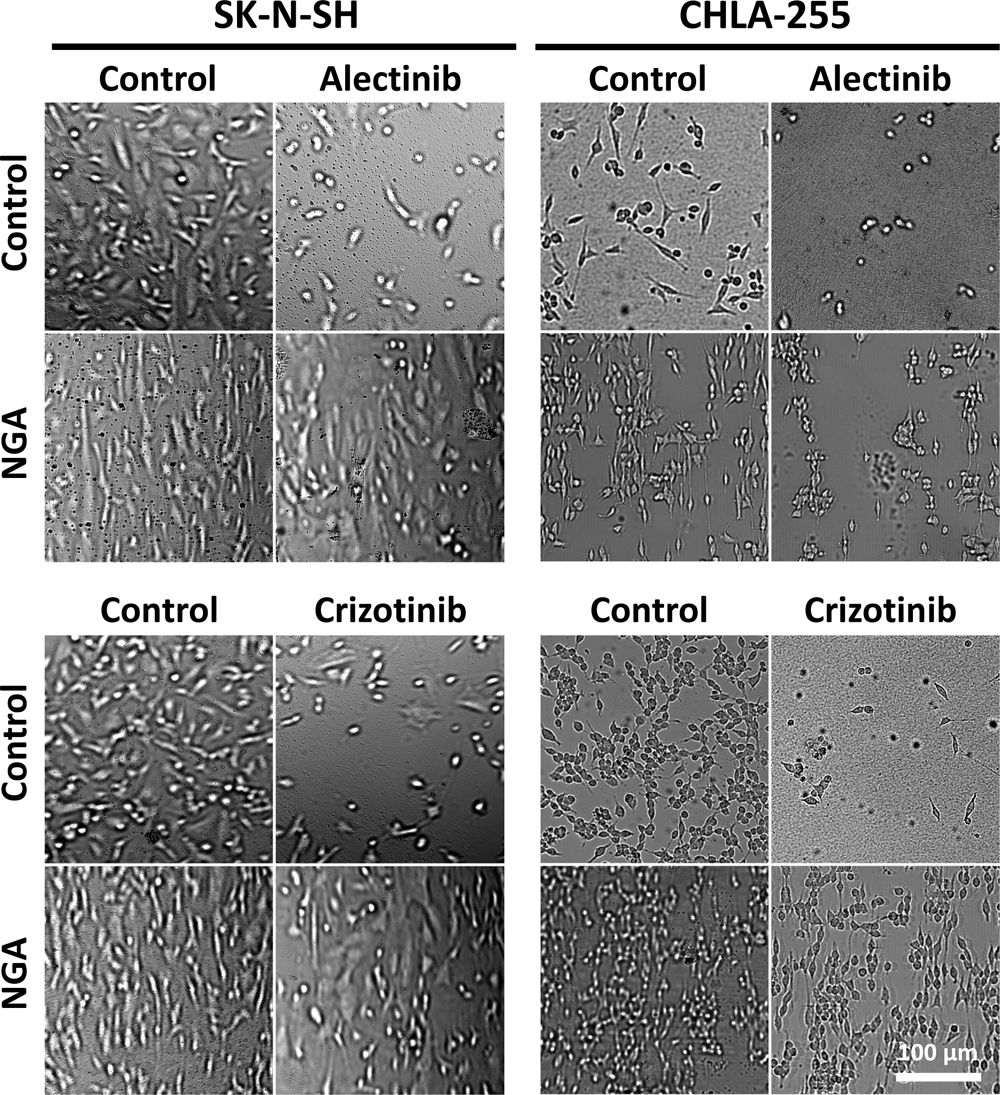
SK-N-SH and CHLA-255 cells on NGA exhibit ALK inhibitor resistance. NB cells were plated on Col-III coated NGA and control substrates. Three random fields of view (n=3) in brightfield microscopy were used for cell counting to determine cell viability following 48 hr incubation with the ALK inhibitors, alectinib and crizotinib. Representative fields of view are shown at the highest dose for alectinib (3 μM) and crizotinib (1 μM) tested at 48hr.

**Supplementary Figure 7.**
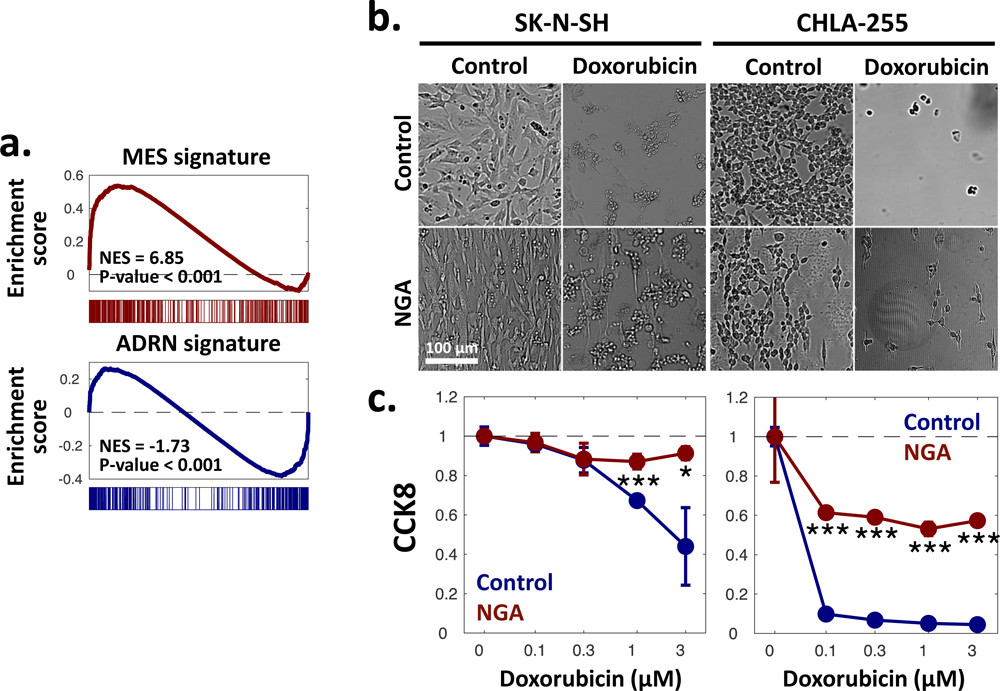
SK-N-SH and CHLA-255 cells on NGA exhibit doxorubicin resistance. **(a)** GSEA analysis of the transcriptome of doxorubicin-resistant NB showing enriched MES and depleted ADRN mRNA signatures **(b)** Three random fields of view (n=3) in brightfield microscopy were used for cell counting to determine cell viability of SK-N-SH and CHLA-255 cells plated on Col-III coated NGA or control substrates following 48 hr incubation with the chemotherapy drug doxorubicin (topoisomerase II inhibitor). Representative fields of view are shown at the highest dose for doxorubicin (3 μM) tested at 48 hr. **(c)** CCK-8 colorimetric assay to quantitatively evaluate cell viability to doxorubicin on Col-III coated NGA versus control substrates after 48hr across a range of drug concentrations. The dots represent the mean value (biological triplicates) and the error bars are SEM. Two-sided Student’s t-test. *P<0.05, **<0.01, ***P<0.005, and ****P<0.001.

**Supplementary Figure 8.**
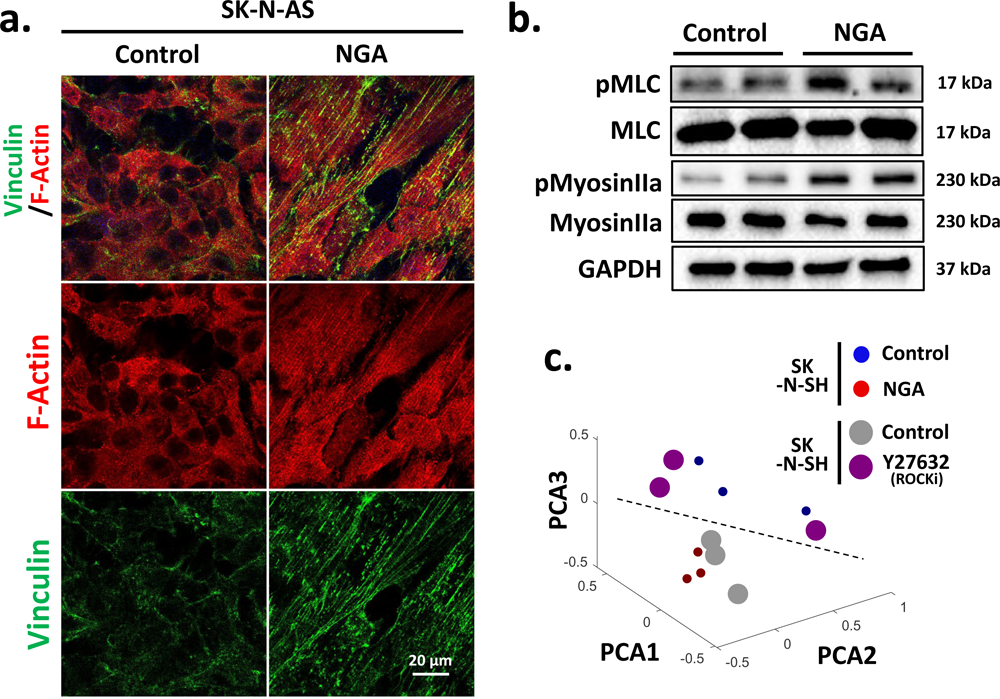
NB cells on NGA display enhanced cell-ECM interactions, cytoskeletal remodeling, and elevated ROCK activity. **(a)** IF staining of SK-N-AS cells on Col-III coated NGA and control substrates against (phalloidin) and vinculin. **(b)** WB of NB cells on Col-III coated NGA and control substrates indicating higher phospho-myosin light chain activity and phospho-myosin II, both are downstream effectors of ROCK activation. GAPDH serves as the loading control **(c)** PCA showing clustering of the transcriptome of SK-N-SH cells plated on control, NGA, and NGA plus ROCKi Y27632 (3 μM).

**Supplementary Figure 9.**
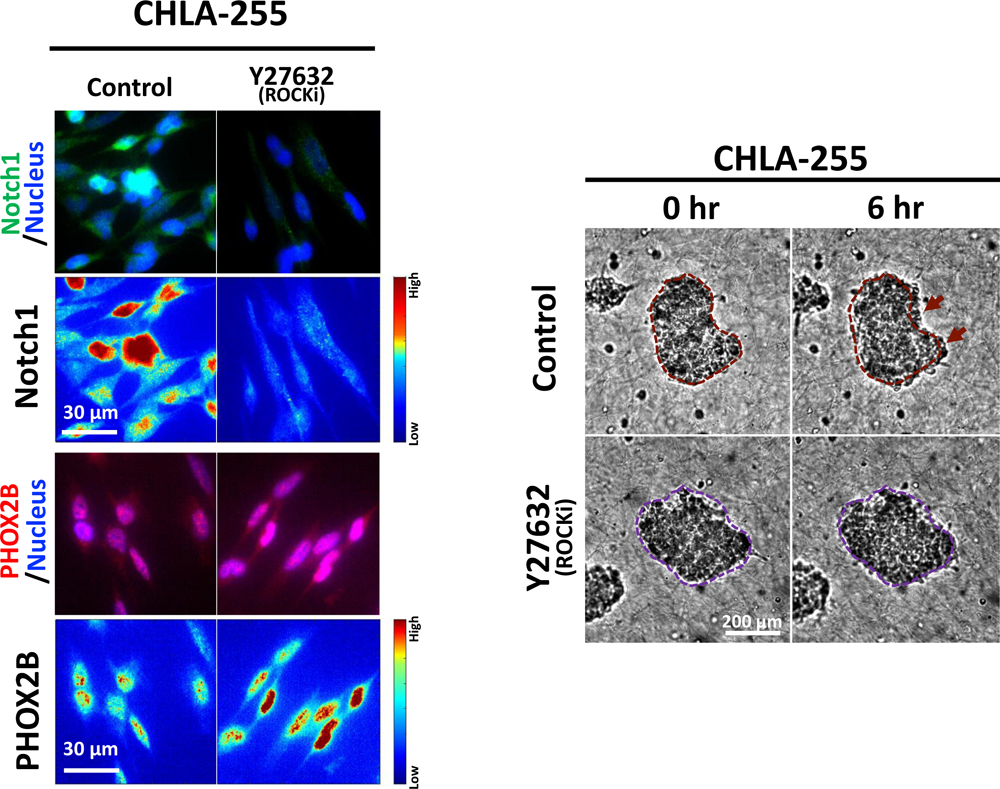
ROCK inhibition abolishes the effects of NGA on the induction of MES markers and invasiveness. **(a)** IF staining of CHLA-255 against PHOX2B (ADRN marker) and Notch1 (MES marker) when plated on NGA (serving as control) versus NGA with ROCKi Y27632 (3μM). Pseudo-color images indicate IF intensity showing higher nuclear levels of PHOX2B and lower nuclear levels of NOTCH1 on NGA with ROCKi Y27632 treatment relative to NGA alone **(b)** CHLA-255 cells were pre-trained on NGA or NGA with Y27632 treatment (3μM) for 3 days and subsequently grown in spheroids via the hanging drop method. NGA and NGA+ ROCKi-trained spheroids were embedded in 3D collagen gels and their expansion dynamics were monitored with time-lapse imaging for 6 hr. White and red dashed lines represent the relative size of spheroids at t=0 and t=6hr respectively

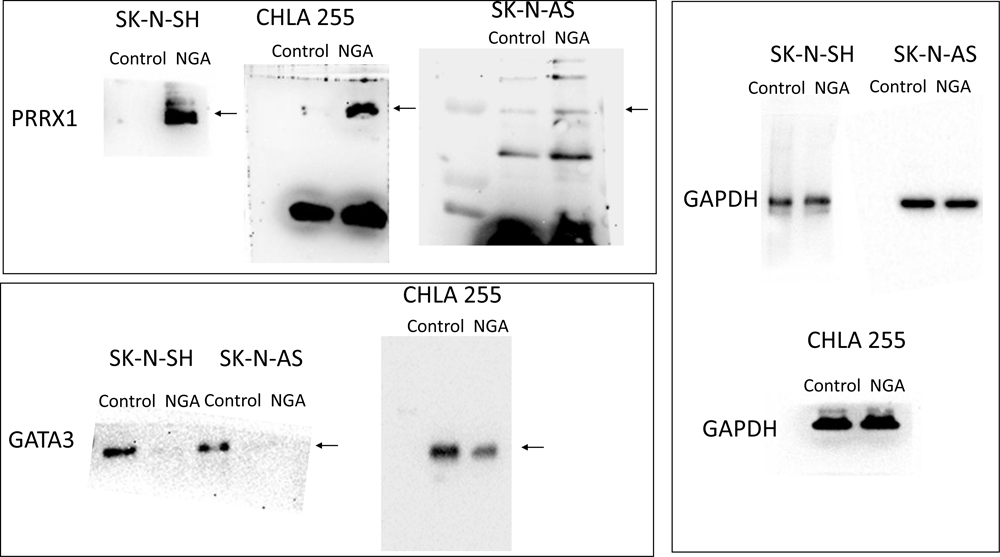

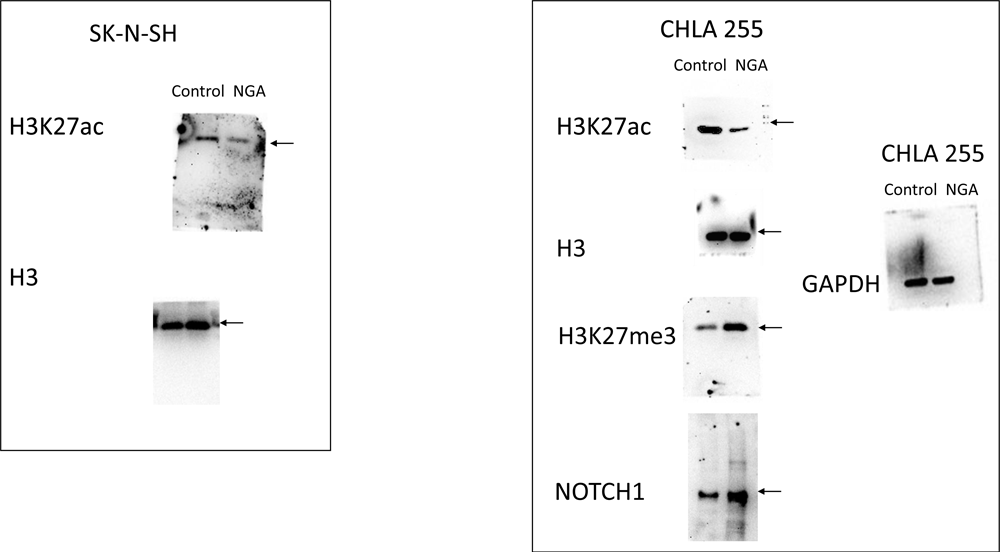

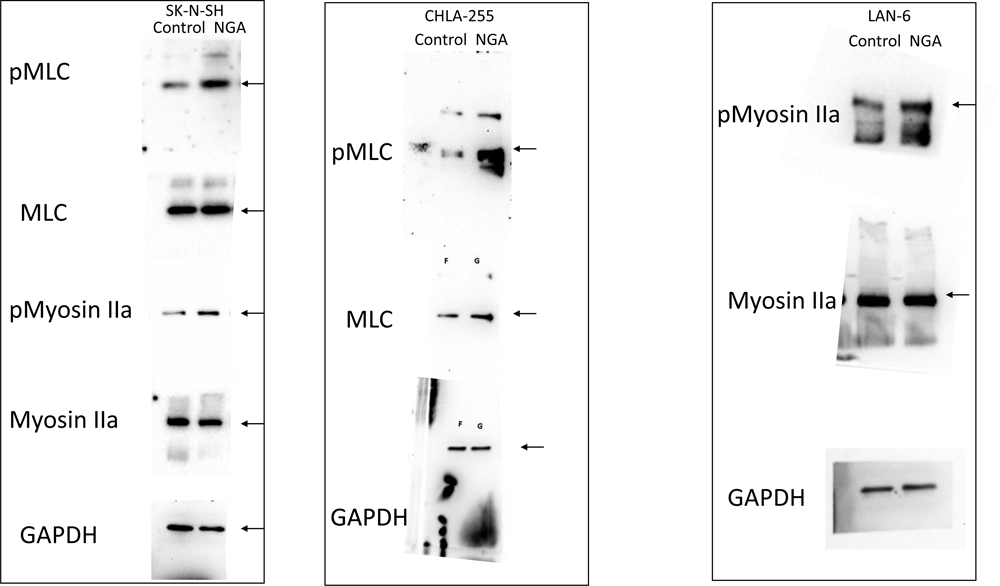

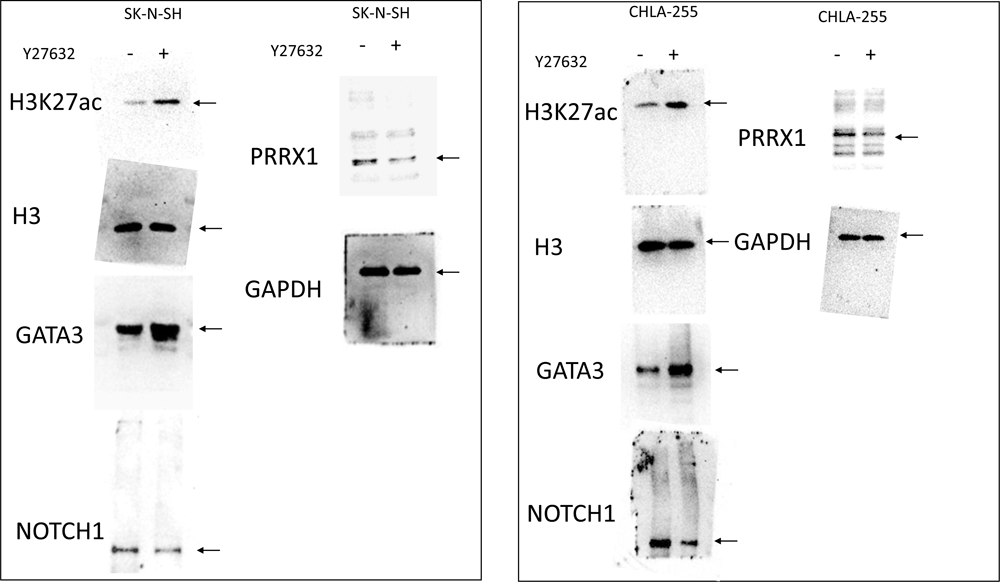

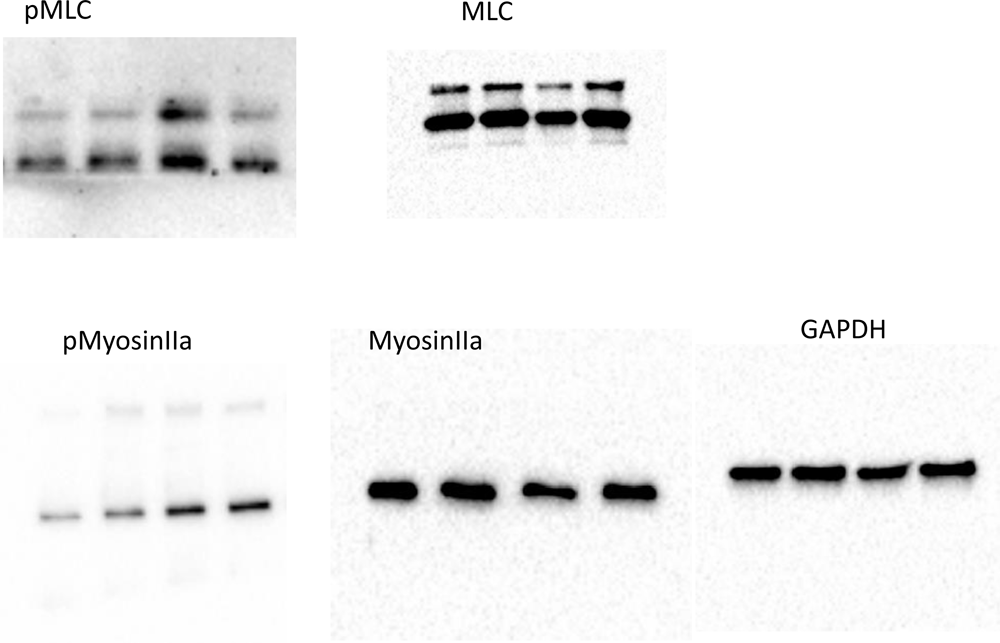

